# More accuracy estimation of the worm burden in the ascariasis of children in Kinshasa

**DOI:** 10.1101/537126

**Authors:** Guyguy Kabundi Tshima, Paul Madishala Mulumba

**Affiliations:** Service de Parasitologie, Département de Médecine Tropicale, Maladies Infectieuses et Parasitaires, Faculté de Médecine, Université de Kinshasa, Democratic Republic of the Congo; Alumnus of the Interuniversity diploma on HIV and AIDS of the Ouagadougou University; Alumnus of the Faculty of Bioscience Engineering of the Ghent University; Free Student at the University of Montreal

**Keywords:** KEY-WORDS, *Ascaris lumbricoides*, estimation of the parasite mass, Kinshasa

## Abstract

The present study aims to give a better estimate of the worm burden (ascariasis) to address accurately the impact of intestinal parasitosis on the children growth in Africa.

The study was conducted on 20 subjects aged 10 months to 10 years (Mean ± SD: 5.6 ± 2.3 years). They were treated with 10 mg/kg of Pyrantel pamoate. The next day, the stools were collected, washed and filtered to harvest all adult ascaris. In total, 141 ascaris (71 males and 70 females) were extracted for 879.9 g of stool. The geometric mean of eggs counted was 29 by 2 mg of stool. The daily eggs laying per female was estimated to 202,500 eggs/days (CI95%: 128,800 – 276,200).

Statistical analysis shows that the parasitic worm burden was proportional both to the number eggs counted per unit of stool volume, and to age of infested subject. A regression model based on these two parameters, with a coefficient of determination equal to 59 %, was retained. Thus, for an old subject respectively of 1, 5 and 10 years, at which 1 egg of ascaris in approximately 2 mg of a preparation would lodge a respective parasitic mass of 1, 3 and 9 g. The results are in the form of confidence interval. For example, for a 5 years old subject with an average of 10 eggs (CI95% = 5.6 - 14.4) after reading of 2 separated preparations coming from the same specimen, the estimated parasitic load is laying between 7 and 11 g.

## INTRODUCTION

The insufficient food resources that still arrive on the dish of the Congolese children are most of the time spoiled by intestinal parasites that they may have had the misfortune to be infected. Thus, with an equal number of parasites infestation, the weight loss will be even greater in those who carry a greater number of *Ascaris lumbricoides*, which is presented as the largest of the intestinal nematodes ^1,2^. It is estimated that an adult *Ascaris lumbricoides* daily spoiled between 1 to 2.8 g of protein per kg of body weight ^1^.

Estimating intestinal parasite load is therefore important in assessing the impact of parasitic diseases on the growth of children in the Democratic Republic of the Congo. To date, we have used the Vandepitte^3^ popularized correspondence scale to relate the number of adult worms in the gut and the number of eggs observed in a stool fragment examined microscopically. According to this author, 2 eggs observed in 2 mg of preparation correspond to an adult worm. The conditions under which this relationship was established are not indicated. That is why the present study set itself the goal of re-evaluating this relationship in our usual working conditions.

## MATERIAL AND METHODS

### 1. Subjects of study

Children aged 10 months to 10 years were selected from the «Centre de Santé Kimbanguiste de kasa-Vubu» and «Centre Ekolo ya Bondeko de Bondo» both institutions are in the Democratic Republic of the Congo.

### 2. Ethical statement

The Department of Tropical Medicine of the University of Kinshasa approved the study. All parents of any child participant provided informed written consent on the child’s behalf.

### 3. Case selection

Direct microscopic stool examination was performed on a standard preparation of approximately 2 mg. The number of eggs found in each case was noted.

The parasitized subjects received the same day a deworming treatment (Pyrantel pamoate) at a rate of 20 mg per kg of body weight. A bottle with a large opening of about 500 ml of capacity was given to them for the collection of all their stools issued within 24 to 48 hours. In total, we collected 20 stool samples containing adult worms. These specimens were brought to the laboratory where they were weighed and sieved through a wire mesh for roundworm harvesting. These, after draining, were weighed in turn using an electronic balance (Metler ®). For each case, the number of male and female worms and their weights were noted.

### 4. Statistical analyzes

The number of worms and their weight relative to the number of eggs or the age of the subjects were adjusted by a linear regression model. The logarithmic transformation has been applied where appropriate to certain variables, such as the weight of the worms and the number of eggs, in order to find a better fit. The quality of the adjustment obtained was evaluated at the probability level of 5% using the coefficient of determination R². The latter represents the amount of information correctly translated by the mathematical model used for the adjustment of the data. The model chosen was the one that gave the largest R². The confidence interval was determined according to Poisson’s law from two measurements made on two preparations made on the same specimen. The daily spawning estimate was based on the geometric mean number of eggs observed in a standard stool volume examined. According to Sinniah B et al ^4^, adult female roundworms start laying when they reach a critical weight of 1.1 g.

The analyzes were performed using SPSS software version 10.0.7 and additional calculations using MathCad version 7 software.

## RESULTS

### 1. Profile of results

On average, we counted 55.5 eggs per preparation and collected 141 worms. These had a sex ratio of 1.01: 1 (71 males and 70 females). Females were significantly larger than males (mean weight: 3.7 1.8 g versus 1.5 0.7 g, mean size: 21.2 4.8 cm vs 15.6 3.0 cm). The extreme values observed were, for the size, between 120 and 320 mm for females as against 102 and 240 mm for males, and for weight, between 0.3 and 7.2 g for females, and between 0.2 and 4.3 g for males. Ten percent of females had a weight of less than 1.1 g as a critical threshold for starting spawning.

The geometric mean number of eggs per preparation was 29. The other average parameters are shown in Table 1.

**Table 1.** Age of study subjects (in years), number of eggs per 2 mg of stool examined the previous day, stool weight (in g), worm weight and number of male and female worms.

| N° | AGE | EGGS | WEIGHT |  | NUMBER |  |  |
| --- | --- | --- | --- | --- | --- | --- | --- |
|  |  |  | STOOL | WORM | WORM | FEMALES | MALES |
| 1 | 3 | 78 | 57,2 | 7 | 2 | 1 | 1 |
| 2 | 6 | 51 | 23,0 | 33 | 6 | 2 | 4 |
| 3 | 0,8 | 11 | 51,2 | 6 | 2 | 1 | 1 |
| 4 | 4 | 4 | 18,7 | 2 | 1 | 1 | 0 |
| 5 | 4 | 10 | 36,4 | 6 | 2 | 1 | 1 |
| 6 | 5 | 63 | 64,0 | 15 | 2 | 1 | 1 |
| 7 | 6 | 96 | 48,4 | 62 | 18 | 8 | 10 |
| 8 | 3 | 15 | 10,7 | 6 | 2 | 0 | 2 |
| 9 | 3 | 257 | 46,8 | 14 | 2 | 1 | 1 |
| 10 | 3 | 10 | 23,0 | 2 | 2 | 0 | 2 |
| 11 | 7 | 8 | 65,4 | 7 | 4 | 2 | 2 |
| 12 | 4 | 76 | 28,7 | 19 | 13 | 5 | 8 |
| 13 | 8 | 30 | 34,9 | 29 | 13 | 7 | 6 |
| 14 | 10 | 6 | 69,0 | 12 | 6 | 2 | 4 |
| 15 | 8 | 177 | 70,7 | 49 | 16 | 10 | 6 |
| 16 | 6 | 8 | 61,1 | 35 | 9 | 6 | 3 |
| 17 | 9 | 13 | 35,9 | 13 | 7 | 4 | 3 |
| 18 | 7 | 75 | 43,6 | 39 | 15 | 7 | 8 |
| 19 | 3 | 49 | 24,3 | 24 | 8 | 4 | 4 |
| 20 | 8 | 28 | 66,9 | 44 | 11 | 7 | 4 |
| TOT | 107,8 | 1065 | 879,9 | 424 | 141 | 70 | 71 |
| MEAN | 5,4 | 53,3 | 44,0 | 21,2 | 7,1 | 3,5 | 3,6 |
| SD | 2,5 | 64,4 | 18,8 | 17,4 | 5,5 | 3,1 | 2,8 |

Legend: TOT: total ; SD: Standard Deviation

Since the examined stool mass was 879.9 g, spawning per female was estimated at 182,300 eggs / day (95% CI = 128,800 to 276,200).

### 2. Adjustment of the data by a linear regression model

There is a significant relationship between the number and weight of parasites depending on either the age of the subjects or the number of eggs counted per volume of their stool (Figures 1, 2 and 3). The relationship between the number of worms and the number of eggs was mediocre (R² = 0.079, p = 0.236).

**Figure 1.**
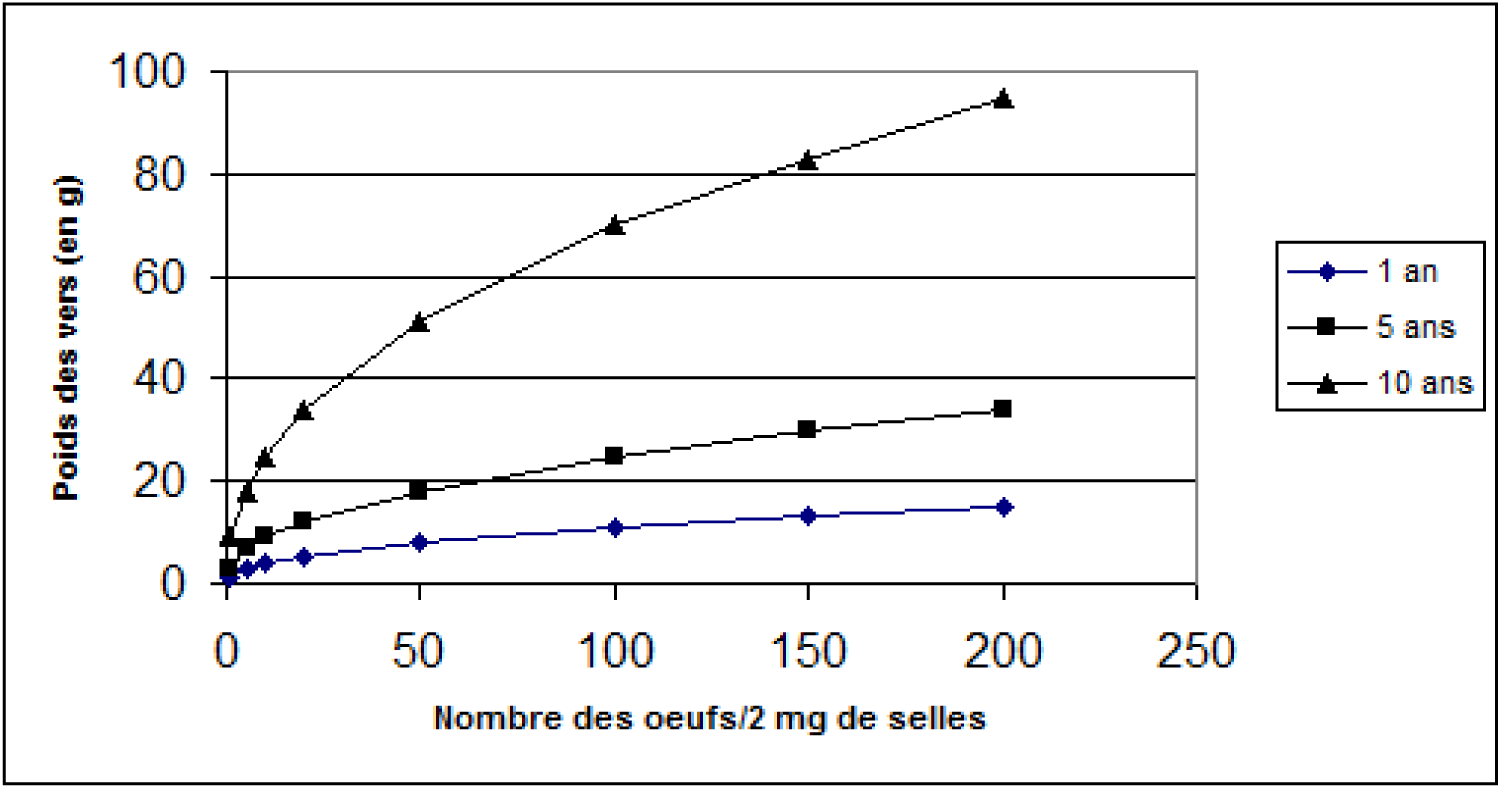
Relationship between the parasite mass (in g), the number of eggs counted in a standard preparation of about 2 mg, and the age of the subjects (in years).

**Figure 2.**
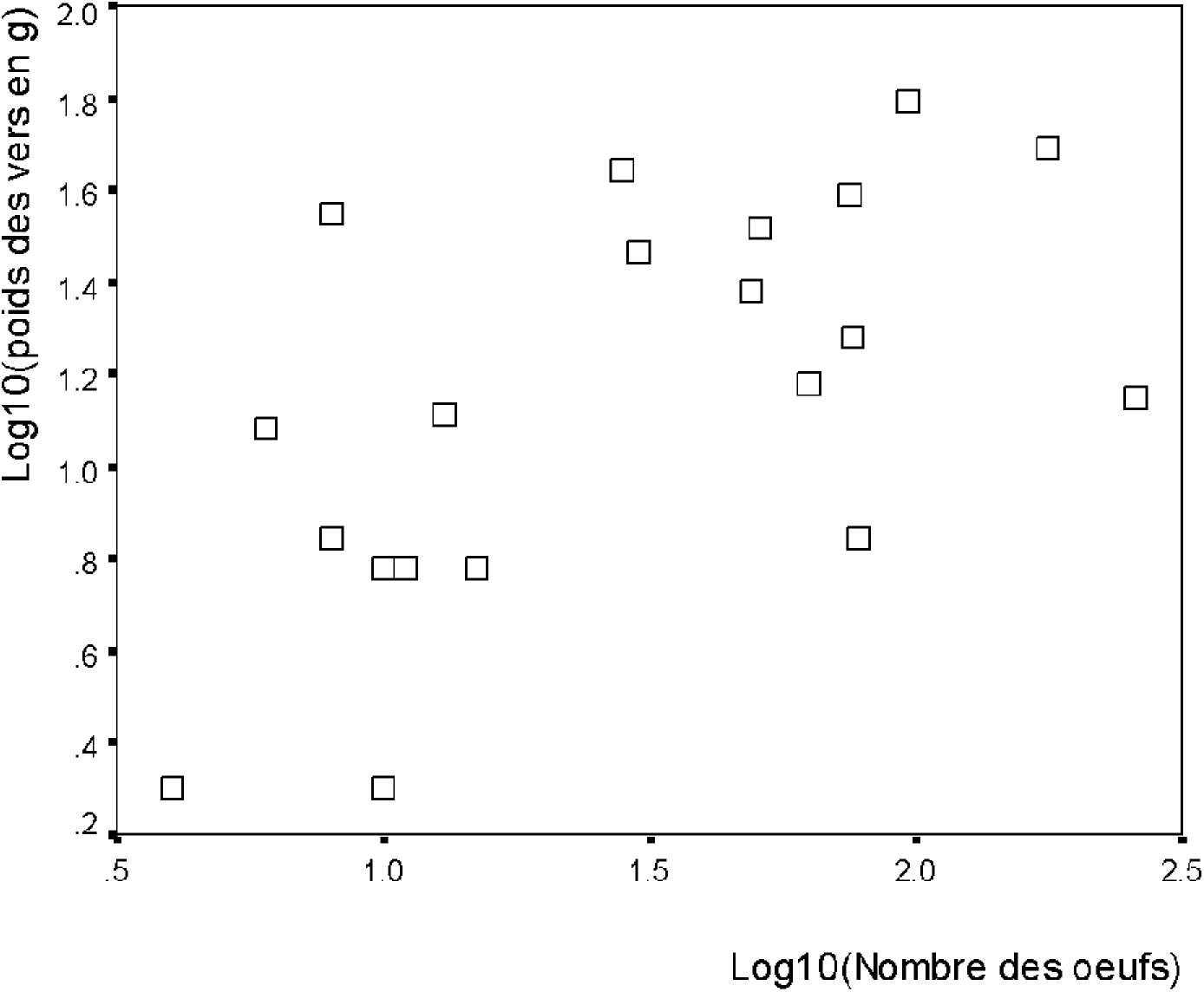
Relationship between worm weight and the number of eggs counted in 2 mg of stool.

**Figure 3.**
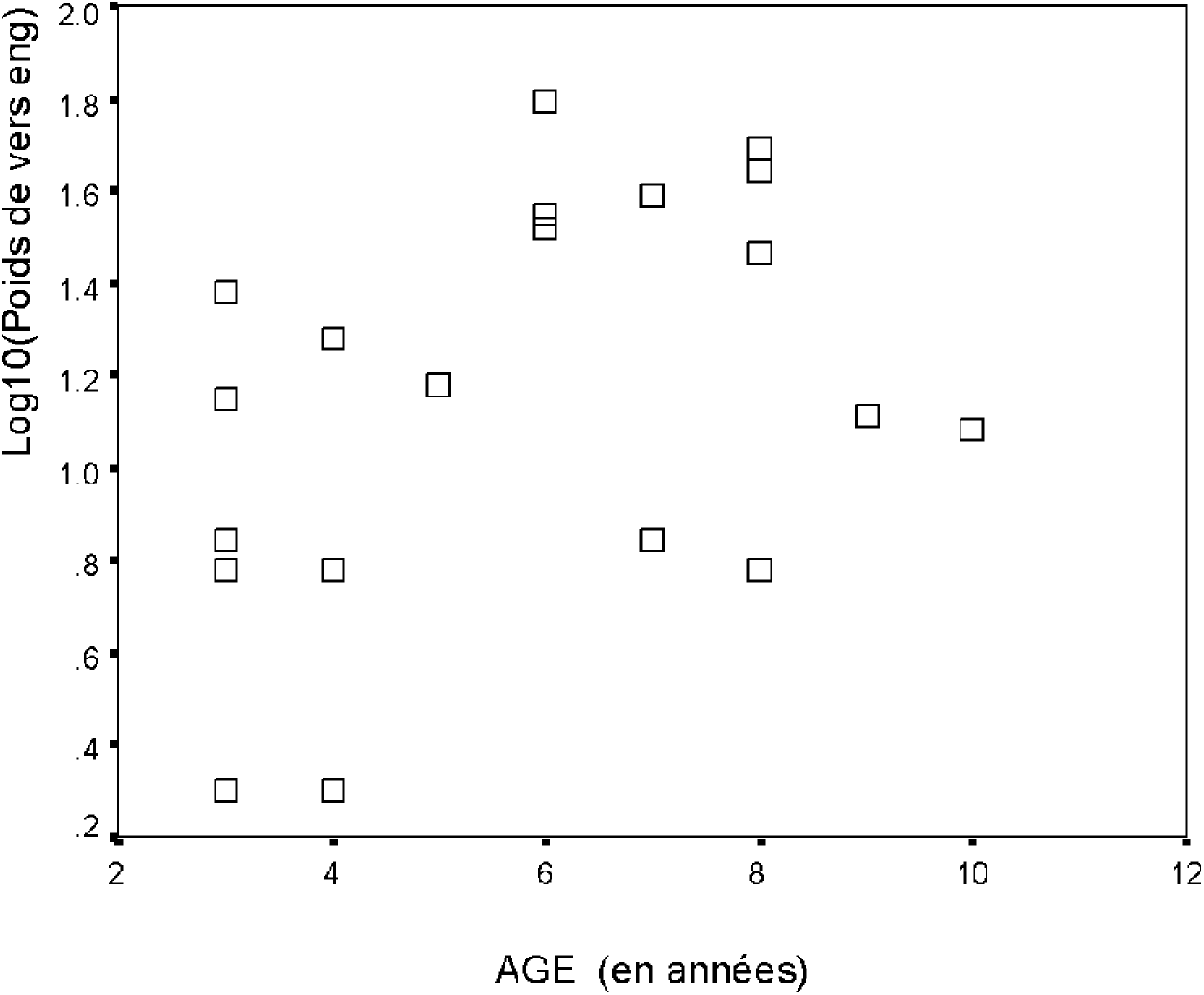
Relationship between age of subjects and number of eggs counted in 2 mg of stool.

**Figure 4.**
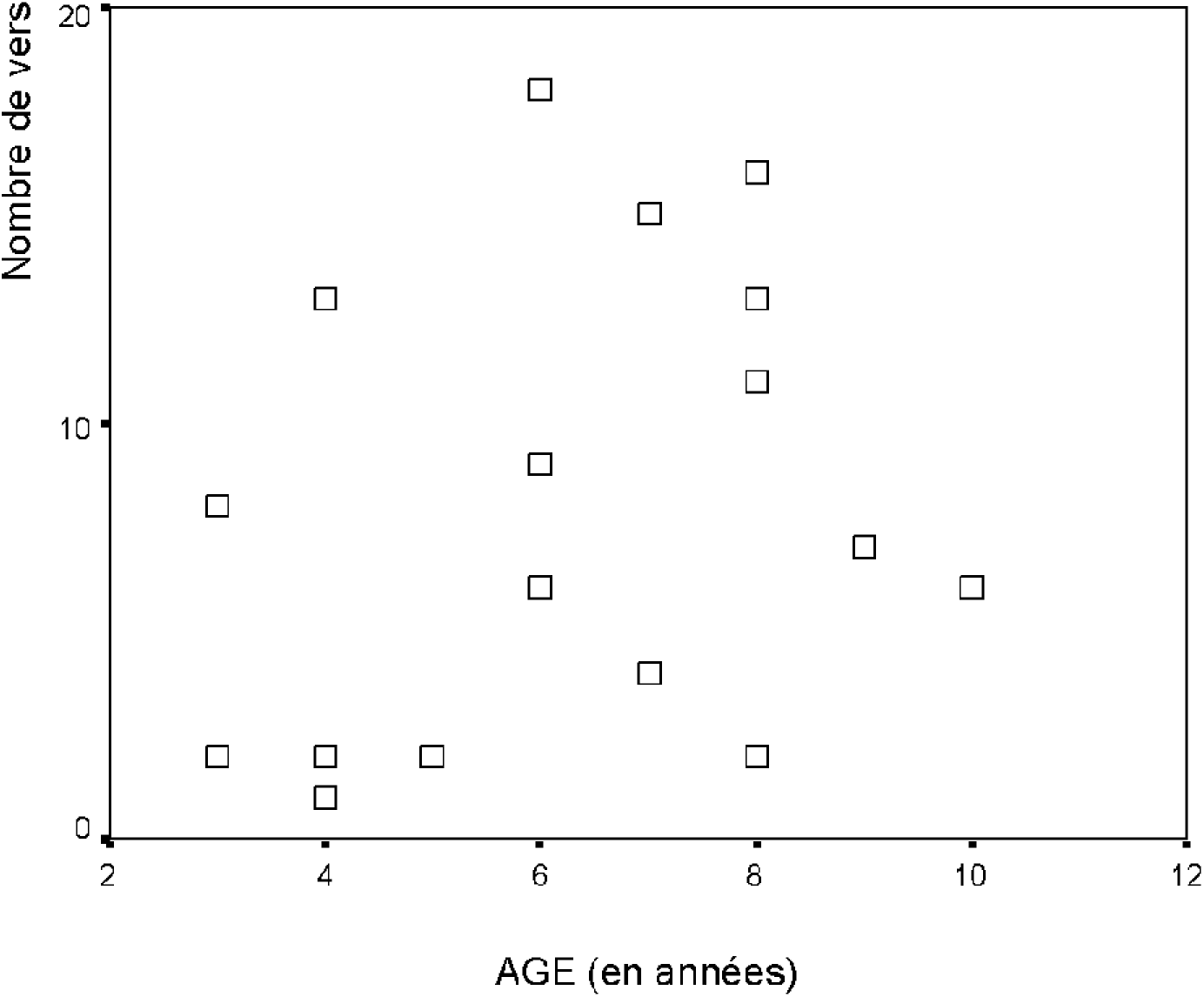
Relationship between age of subjects and number of eggs counted in 2 mg of stool.

**Figure 5.**
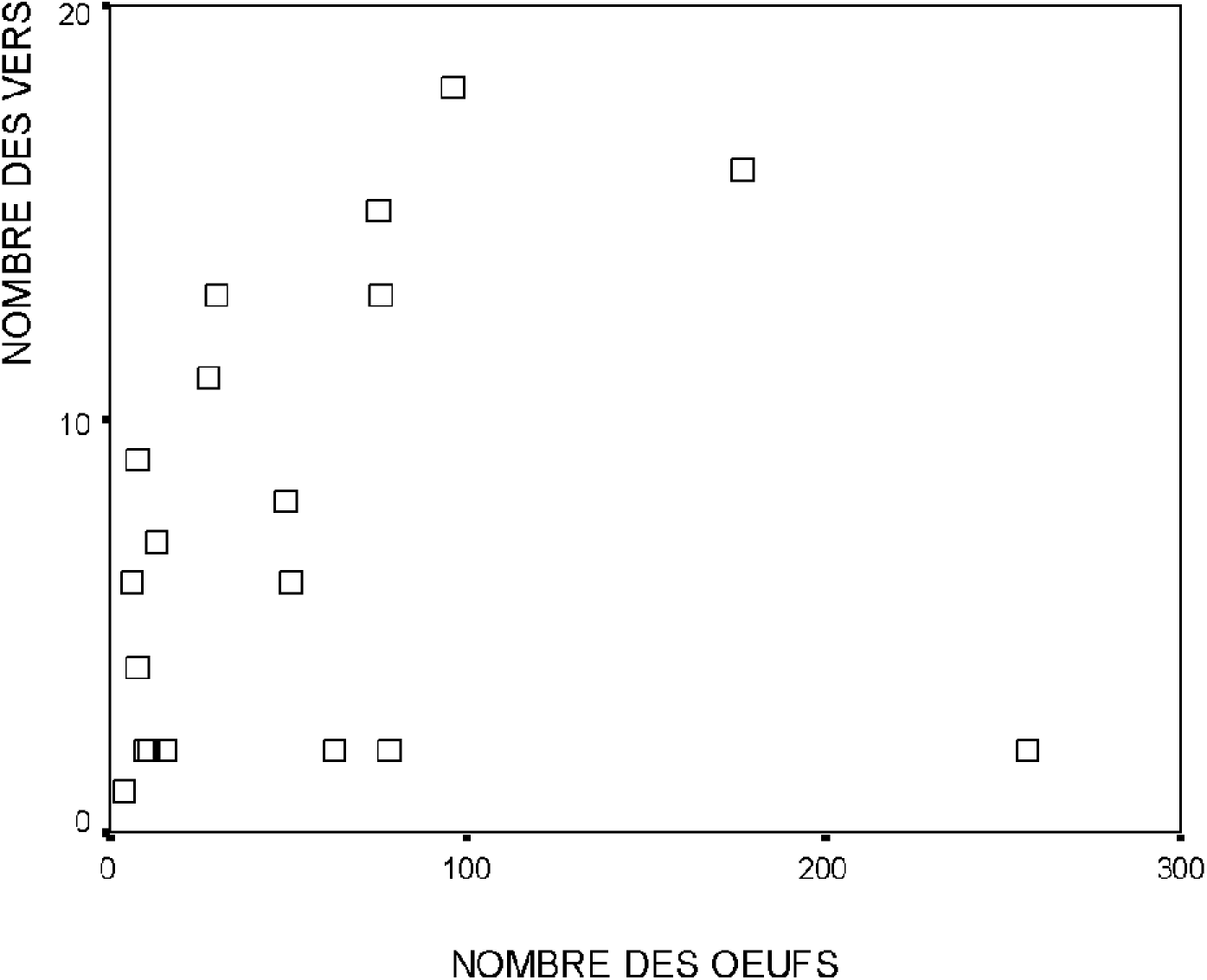
Relationship between the number of worms and the number of eggs counted in 2 mg of stool

The best retained linear regression model, which had a coefficient of determination (R²) equal to 0.590 (p = 0.001), the highest of all, which predicted age and number of eggs; had the following form:

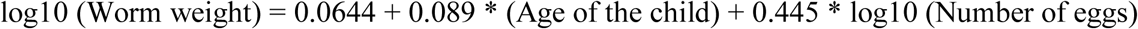

It should be noted that the regression found between worm weights and each of the individual variables was relatively less effective. In fact, the value of R² obtained with the age of the subjects and the number of eggs was respectively equal to 0.253 (p = 0.024) and 0.296 (p = 0.013).

As an illustration, from the model chosen and for a standard preparation of 2 mg containing 1 egg of *A. lumbricoides*, the parasite mass for a subject aged respectively 1, 5 or 10 years will be 1, 3 and 9 g.

### 3. Confidence interval of the results

For example, for a 5-year-old child with an average of 10 eggs (95% CI = 5.617-14.383) after reading 2 independent preparations, the estimated parasite load will be between 7 and 11 g. The limits of this confidence interval were calculated according to the parameters of the regression model used:

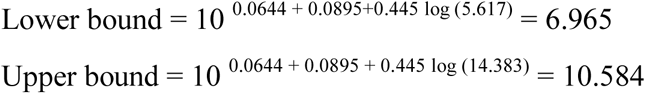

This interval depends on the 95% CI of the number of eggs obtained according to the Poisson’s law:

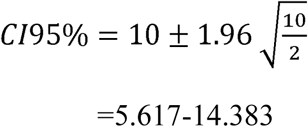

## DISCUSSION

A smear containing two eggs, according to Vandepitte ^3^, corresponds to an adult roundworm. This last result seems to us to come from a simple arithmetic deduction. Indeed, since the adult female lays 200 000 eggs / day and the faecal bowl weighs about 100 g. In this case, 2 eggs found in 2 mg of stool correspond to 100,000, which is half of the spawning of a female. If we accept the hypothesis of parity between the number of males and females, this number must be multiplied by 2, which amounts to an adult.

In the present study, the relationship we found between the number of worms or their mass relative to the number of eggs observed in a standard stool fragment is far from simple. On the contrary, our results showed that the parasite mass, better than the number of worms, was significantly related to the number of eggs counted per unit of stool volume (Figure 1) and the age of the subjects (Figure 2). In fact, the oldest subjects presented the largest worms compared to younger subjects. Thus, this result suggests that the influence of age can be considered in estimating the parasite load.

Not only the number of eggs found, but also the number of gravid females present in the digestive tract, varies from one subject to another, and the number of eggs laid per female can show significant circadian or seasonal variations. In fact, according to Sinniah B et al ^4^, adult female roundworms begin to lay when they reach the size of 118 mm for a diameter of at least 3.7 mm. The average minimum weight of a female producing eggs is 1.1 g and would also depend on the age of the female. In this study, 10% of females had a weight less than this value. In addition, congestion of the intestinal microenvironment due to the massive presence of parasites has a negative impact on oviposition ^5^. In other words, the law of diminishing returns reflects the effect of the reduction of individual food resources in the face of an inconsiderate increase in population. Other more complex interactions are likely to influence egg laying parameters, such as the abundance and composition of the bacterial flora, the presence of other parasites, host immunity and histocompatibility. On this last point, blood group A is particularly thought of. In fact, subjects in this blood group are more susceptible to *A. lumbricoides* infections and have higher parasite mass than the rest of the population ^6^.

In addition, according to the work of Sinniah et al ^4,6,7^, the average daily spawning is estimated at 238,722 (95% CI = 134,462 to 358,750). Our results are contained in the confidence interval found by these authors. The sex ratio found by Sinniah et al 1 male against 1.4 female corroborates somewhat that found in the present work which was 1.01 male against 1 female, in both cases, close to parity.

Given the importance of the hazards of spawning, for a better estimate of the parasite mass from this parameter, we recommend carrying out at least two independent measurements from which the mean and the confidence interval will be calculated, preference is given to the geometric mean to minimize the amplitude of fluctuations between observations ^8^.

In conclusion, estimating the parasite mass of *A. lumbricoides* is a complex undertaking that needs further refinement. We believe that while estimating the parasite mass for a given individual remains quite risky, it is very valuable for estimating average parasite mass in community surveys. To improve the quality of our estimates, we must increase the size of our samples, on the one hand, and integrate the possible influence of the environment, on the other hand.

